# Intra-Lymph Node Crosslinking of Antigen-Bearing Polymers Enhances Humoral Immunity and Dendritic Cell Activation

**DOI:** 10.1101/2024.03.13.584831

**Authors:** Erin M. Euliano, Anushka Agrawal, Marina H. Yu, Tyler P. Graf, Emily M. Henrich, Alyssa A. Kunkel, Kevin J. McHugh

## Abstract

Lymph node (LN)-resident dendritic cells (DCs) are a promising target for vaccination given their professional antigen-presenting capabilities and proximity to a high concentration of immune cells. Direct intra-LN injection has been shown to greatly enhance the immune response to vaccine antigens compared to traditional intramuscular injection but is infeasible to implement clinically. Employing the passive lymphatic flow of antigens to target LNs has been shown to increase total antigen uptake by DCs more than inflammatory adjuvants, which recruit peripheral DCs. Herein, we describe a novel vaccination platform in which two complementary multi-arm poly(ethylene glycol) (PEG) polymers—one covalently bound to the model antigen ovalbumin (OVA)—are injected subcutaneously into two distinct sites that drain to the same LN through different lymphatic vessels and, upon meeting in the LN, rapidly crosslink. This system improves OVA delivery to, and residence time within, the draining LN compared to all control groups, with the crosslinking of the two PEG components improving humoral immunity without the need for any pathogen-mimicking adjuvants. Further, we observed a significant increase in non-B/T lymphocytes in LNs cross-presenting the OVA peptide SIINFEKL on MHC I over a dose-matched control containing alum, the most common clinical adjuvant, as well as an increase in DC activation in the LN. These data suggest that this platform can be used to deliver antigens to LN-resident immune cells to produce a stronger humoral and cellular immune response over materials-matched controls without the use of traditional adjuvants.

**Translational Impact Statement:** Vaccines save millions of lives each year; however, they often require more than one injection to confer protection and do not always provide long-term immunity. Transitioning from intramuscular injections to intra-lymph node injections has been shown to greatly increase immunity conferred by vaccination but is infeasible to implement clinically. Herein, we present a biomaterial-based vaccination strategy that delivers antigen to the lymph nodes after a pair of remote injections, which increases the duration of immune cell exposure to the vaccine and, consequently, enhances humoral immunity.

## Introduction

Vaccination is an effective strategy for reducing the spread of infectious diseases. Most current vaccines are administered in a series of intramuscular injections over the course of weeks or months. After injection, the antigen either passively drains to a regional lymph node (LN) or is taken up by an antigen-presenting cell (APC) and carried to the LN, where it briefly interacts with lymphocytes before being cleared.^1^ This short-lived interaction between antigens and B and T cells often generates only a marginal immune response after one dose that leaves many individuals still at risk of contracting the disease or protected for only a short duration.^2^ Therefore, most vaccines require the use of multiple doses delivered at optimized intervals to induce secondary immune responses that confer immunity through memory B cell and plasma cell production, antibody affinity enhancement, and/or T cell expansion and differentiation. However, there are cases in which an optimized schedule provides only limited protection; for the malaria vaccine recently endorsed by the World Health Organization, RTS,S/AS01, the four recommended injections provide protection for only 36% of children^3^ and the vaccine has several potential side effects.^4,5^ Similarly, GlaxoSmithKline’s tuberculosis vaccine, M72/AS01E, provides less than 50% protection after the recommended two doses.^6^ Although the development of better antigens and/or adjuvants could increase these relatively low response rates, identifying such candidates has proven challenging for these and many other diseases. The reasons for this include a lack of defined immunological correlates of infection, the evasion of host immunity by the pathogen, and a limited understanding of the pathogen itself.^7,8^ Alternative strategies that better control the spatial and temporal interactions between lymphocytes and antigens may have the ability to eliminate the need for multiple clinical visits, enhance seroconversion rates, extend the duration of immunity, and/or rescue previously non-viable vaccines.

Extending the release of vaccine antigens has been shown to produce stronger immune responses as compared to bolus doses,^9–11^ as has increasing the residence time of antigen within the LN.^12,13^ Similarly, direct antigen injection into LNs has been shown to vastly improve vaccine immunogenicity, but is infeasible at scale due to the need for imaging technology to precisely guide needle placement and the consequences of an off-target injection.^14^ Various nanoparticle carriers have been used to target vaccines to the LN and, while this strategy has generally increased LN residence time, the improvement is typically marginal—on the order of 24-48 h.^13,15–19^ The particle size required to drain into the lymphatics from the subcutaneous space has been well-studied;^20,21^ it is generally reported that carriers must be at least 5-10 nm in diameter to minimize uptake into blood capillaries but less than 100 nm to maintain mobility within the extracellular matrix.

LN-resident dendritic cells (DCs), which are predominantly immature, have been shown to take up antigen and activate T cells entirely within the LN if antigen is co-delivered with a maturation stimulus.^22^ Additionally, the efficient targeting of antigen to the LN has been shown to induce a higher percentage of antigen-presenting DCs compared to traditional strategies that target migratory peripheral DCs using adjuvants like aluminum hydroxide (alum).^2^ Targeting antigen to the LN via passive lymphatic flow creates an opportunity to avoid the use of locally inflammatory adjuvants, thereby reducing potential side effects.^23^ We present a novel vaccine delivery strategy that enhances LN accumulation and residence time of a model antigen, ovalbumin (OVA), through the rapid crosslinking of two complementary polymers within the LN via a bioorthogonal click reaction^24^ following their passive passage through different lymphatic vessels. To ensure the safety and reliability of the crosslinking reaction, we employed the reaction between trans-cyclooctene (TCO) and tetrazine (Tz), an inverse-electron demand Diels-Alder cycloaddition, which is the fastest bioorthogonal click reaction identified to date.^25,26^ This enabled the components to crosslink rapidly even at the low concentrations present in the LN, resulting in an improved humoral response and increased DC activation over dose-matched controls.

## Materials and Methods

### Materials Synthesis and Characterization

Eight-arm, 40 kDa amine-reactive PEG (succinimidyl succinimide ester, SAS) (Creative PEGWorks, Durham, NC) was reacted with trans-cyclooctene (TCO)-amine or Tetrazine (Tz)-amine (Vector Laboratories, Newark, CA) in dichloromethane (DCM). TCO-PEG conjugated to Cyanine5-amine dye (BroadPharm, San Diego, CA) was made by reacting 1:12:0.2 molar equivalents PEG:TCO-amine:Cy5-amine in DCM containing 5 equiv. DIEA overnight at room temperature (RT), protected from light. Tz-PEG was synthesized by reacting PEG with Tz-amine at a molar ratio of 1:12 in DCM containing 5 equiv. DIEA overnight at room temperature, optimized to ensure that approximately 1 out of 8 PEG arms retained an amine-reactive SAS group capable of attaching OVA in a subsequent reaction.

Both PEG solutions were dried using a Genevac Centrifugal Evaporator (Scientific Products, Warminster, PA), resuspended in Milli-Q water, and filtered through 100 kDa MWCO Amicon Spin Filters (MilliporeSigma, Burlington, MA) to remove any particulates. Solutions were then purified using 3 kDa MWCO Amicon Spin Filters to remove low molecular weight TCO-amine or Tz-amine. Finally, the materials were passed through PD-10 Desalting Columns (Cytiva, Marlborough, MA) to remove any remaining reactants.

Ovalbumin was reacted with 0.5 equiv. DyLight 755-NHS Ester (both from Thermo Fisher Scientific, Waltham, MA) in 0.5 M sodium bicarbonate buffer at pH 9.6 for 2 h at room temperature, protected from light. Unreacted fluorophore and residual endotoxin were removed by size exclusion chromatography (SEC) using a HiLoad Superdex 200 pg preparative SEC column (Cytiva). Purified OVA-DyLight 755 was then reacted at a 1:1 molar ratio with Tz-PEG in 0.5 M sodium bicarbonate buffer at pH 9.6 for 2 h at room temperature, protected from light. Given that the remaining amine-reactive NHS groups on Tz-PEG were hydrolytically labile, this step occurred quickly following Tz-PEG purification. SEC was then used to separate Tz-PEG-OVA from unconjugated OVA (Figure S1).

The degree of Tz and TCO conjugation to PEG was calculated using ultraviolet-visible (UV-VIS) spectroscopy. Tz has a maximum absorbance at 530 nm, enabling the degree of labeling to be calculated by comparing the concentration of Tz measured by absorbance to that of PEG measured by mass. TCO does not absorb in the visible light spectrum, but reacting Tz with TCO causes Tz to no longer absorb light at 530 nm. The degree of TCO labeling was therefore calculated by reacting TCO-PEG with an excess of Tz-amine and calculating the concentration of Tz reacted. The degree of labeling of Cy5 on TCO-PEG and of OVA-DyLight 755 on Tz-PEG were calculated by measuring their fluorescence (Ex/Em 630/670 nm and 750/790 nm, respectively) on a microplate reader (Tecan, Männedorf, Switzerland) compared to known standards. The approximate hydrodynamic diameters of the materials were calculated via dynamic light scattering using a Zetasizer Nano (Malvern Panalytical, Worcestershire, UK).

### Endotoxin Testing

All materials for *in vivo* use were tested for endotoxin using the HEK-Blue mTLR4 reporter cell line (InvivoGen, San Diego, CA) following the manufacturer’s protocol. In brief, PEG and OVA materials were dissolved in Milli-Q water at 1 mg/mL and 20 µL of each solution was added per well to a 96-well tissue culture-treated plate. An endotoxin bacterial lipopolysaccharide standard curve was also plated over a range of 0.15-10 endotoxin units (EU) per mL. HEK-Blue mTLR4 cells seeded at a density of 2.2 x 10^5^ cells/mL in 180 µL of treated growth media (DMEM containing 10% v/v FBS, 1% PenStrep, 1x HEK Blue Selection Media, 0.2% Normocin) were then pipetted into each well and incubated at 37 °C and 5% CO_2_ for 20 h. Then, 20 µL of supernatant was added to 180 µL QuantiBlue detection buffer (InvivoGen) and incubated at 37 °C and 5% CO_2_ for 1 h, at which point the absorbance at 630 nm was measured on a microplate reader (Tecan) and endotoxin content was calculated by comparison to the standards.

### *In Vitro* Release

Blind Well Chambers (Neuro Probe, Gaithersburg, MD) with an upper well:lower well capacity of 800:200 µL were used to assess the *in vitro* degradation rate of the hydrogels. Ten µL of each PEG component were mixed at a final concentration of 7.5% (w/v) in the dry lower chamber, then 150 µL of phosphate-buffered saline (PBS) was pipetted on top and the lower chamber was covered with a 13 mm diameter polycarbonate membrane with 3 µm pores (Whatman, Maidstone, UK). The upper chamber was screwed in above the filter membrane and filled with 800 µL PBS, then sealed with a piece of adhesive PCR Seal (Thermo Fisher Scientific). The blind well chambers were attached to a rotisserie spinner and rotated at 37 °C protected from light. Sampling was performed by peeling back the adhesive tape, pipetting out 800 µL from the upper chamber without disturbing the filter, and adding 800 µL of fresh PBS. The fluorescence of the samples was measured on a microplate reader (Tecan) and compared to a standard curve.

### *In Vivo* Administration

All animal studies were conducted in accordance with IACUC-approved protocol 22-246 in the Animal Resource Facility at Rice University in Houston, TX. Female SKH1-Elite mice were purchased from Charles River Laboratories (Wilmington, MA) at 5-6 weeks old and vaccinated at 6-8 weeks old.

All OVA-containing materials were prepared to achieve a final concentration of 1 mg/mL OVA in PBS, resulting in 20 µg OVA in each 20 µL injection. Tz-PEG-OVA at 1 mg/mL OVA contained ∼3.6 mg/mL Tz-PEG, so an equivalent concentration of Tz-PEG was added to unconjugated OVA to constitute the Tz-PEG + free OVA injections. The alum solution was made by matching the amount of PEG by mass to the aluminum in Alhydrogel (alum, InvivoGen), resulting in a solution of 36% alum by volume in PBS. Non-OVA-containing TCO-PEG-Cy5 was dissolved at 75 mg/mL in PBS.

All injections were 20 µL in volume and were performed using 31-gauge insulin syringes (Becton Dickinson, Franklin Lakes, NJ). Mice were anesthetized using a continuous flow of 2.5% isoflurane. Tail base injections were administered by positioning the mouse on its stomach and inserting the needle subcutaneously approximately 0.5 cm to the right of the base of the tail. Ankle injections were administered by positioning the mouse on its back, extending the leg by hand, and inserting the needle just to the inside of the medial malleolus. Mice receiving injections into both the ankle and tail received them within 90 sec of one another.

### *In Vivo* Imaging

SKH1-Elite mice (n=6 per group) were vaccinated as described above and imaged using a PerkinElmer In Vivo Imaging System (IVIS). Animals were anesthetized using a continuous flow of 2.5% isoflurane and placed onto the heated platform in the IVIS chamber. Images were collected using excitation/emission filters at 640/700 nm and 745/800 nm for Cy5 and DyLight 755, respectively. Mice were positioned in the chamber on their side to view the draining inguinal lymph node (InLN). Mice were imaged the day before vaccination, immediately following vaccination, 6 h after vaccination, and each day thereafter for one week. Radiant efficiency in the LN was calculated by positioning an equally-sized circular region of interest over the visible InLN.

### Immunofluorescence of Lymph Node Sections

SKH1-Elite mice (n=3 per group) were vaccinated as described above and euthanized 24 h after injection. Draining InLNs were harvested and placed in formalin overnight at 4 °C. LNs were subsequently moved into 15% sucrose in Milli-Q water until they sank, then transferred to 30% sucrose in Milli-Q water and stored at 4 °C overnight. LNs were then gently dried, excess fat was removed, and the tissue was frozen into Optimal Cutting Temperature (OCT) compound on dry ice. Samples were stored at -80 °C before being cut into 5 µm sections using a cryostat.

Slides containing LN sections were first blocked using SuperBlock Buffer (Thermo Fisher Scientific) for 30 min at room temperature, then treated with a rabbit anti-OVA antibody diluted in SuperBlock Buffer overnight at 4 °C. Slides were rinsed with PBS and treated with fluorescently labeled antibodies (Table S1) diluted in SuperBlock Buffer overnight at 4 °C. Slides were rinsed with PBS and mounted in the dark with Fluoromount-G (Invitrogen, Waltham, MA). Sections were imaged using an EVOS M5000 microscope (Thermo Fisher Scientific).

### Serum Collection and Antibody Quantification

The SKH1-Elite mice used in the IVIS imaging study described above (n=6 per group) were bled by submandibular vein puncture the day before vaccination and 2, 4, 6, 8, 10, and 14 weeks after vaccination in clotting microvettes (Sarstedt, Nümbrecht, Germany). After collection, blood was allowed to clot before being centrifuged at 4 °C for 10 min at 10,000 rcf. The separated serum was collected and stored at -20 °C until use. OVA-specific antibody titers were measured by enzyme-linked immunosorbent assay (ELISA). Nunc MaxiSorp plates (Thermo Fisher Scientific) were coated with 100 µL of 1 µg/mL OVA in pH 9.6 carbonate-bicarbonate buffer and incubated on an orbital shaker overnight at 4 °C. Plates were then washed 3x in 0.5% Tween 20 in PBS (PBST) using a plate washer (BioTek, Winooski, VT), followed by blocking with 300 µL of 5% dry milk (Rockland Immunochemicals, Pottstown, PA) in PBST (blocking solution) for 2 h at room temperature. In secondary non-adhering plates, samples were diluted in blocking solution, beginning at a ten-fold dilution with two-fold dilutions proceeding across the plate. The blocking solution was removed from the MaxiSorp plates and 50 µL of each sample was transferred from the secondary plates to each well. The plates were incubated on a shaker for 2 h at room temperature before being washed 3x with PBST using a plate washer. Next, 100 µL of horseradish peroxidase (HRP)-conjugated rabbit anti-mouse IgG (Jackson ImmnoResearch, West Grove, PA) diluted 1:1000 in blocking solution was added to each well. Plates were incubated on a shaker for 2 h at room temperature, then washed 5x with PBST using a plate washer. Wells were developed using 100 µL SureBlue TMB solution (SeraCare, Milford, MA) and the reaction was stopped after 3 min by adding 100 µL 1N sulfuric acid solution. The absorbance of each well at 450 nm (with a 650 nm reference measurement) was determined using a microplate reader. Antibody titers were reported as the last dilution at which sample absorbance was at least two-fold greater than that of naïve serum.

Antibody subclass titers were measured using serum collected ten weeks after vaccination using the same protocol as described above, though with different secondary antibodies. HRP-conjugated goat anti-mouse IgG1, IgG2a, IgG2b, and IgG3 antibodies (Jackson ImmunoResearch) were each substituted for anti-mouse IgG and diluted 1:1000 in blocking solution. Samples were run at a 25-fold dilution followed by serial two-fold dilutions across the plate. All subclasses were developed for 3 min except for IgG3, which was developed for 10 min.

### Flow Cytometry

SKH1-Elite mice (n=5 per group) were vaccinated as described above. One week after vaccination, mice were euthanized and their spleens and InLNs (both draining and contralateral) were harvested for analysis by flow cytometry. LNs were digested using a protocol modified from the Hubbell group.^27^ In brief, LNs were incubated in 1 mL complete DMEM (+ 1% Pen/Strep and 10% FBS) containing 1 mg/mL collagenase IV and 40 µg/mL DNAse I (both Sigma-Aldrich, St. Louis, MO) at 37 °C for 30 min. Tubes were centrifuged at 250 rcf for 5 min, the supernatant was removed, and 1 mL of complete DMEM containing 2 mg/mL collagenase D (Sigma-Aldrich) and 40 µg/mL DNAse I was added. The tubes were inverted to resuspend pelleted cells and tissue and then incubated on an orbital shaker at 37 °C for 15 min. The tissue was then mashed against the side of the tube using a 1000 µL pipette tip and pipetted approximately 100 times to break up the LN capsule. One mL of ice-cold 10 mM EDTA in PBS was added to quench enzymatic activity and the solution was again pipetted approximately 50 times. Finally, the cell solution was allowed to drip through a 70 µm cell strainer (Fisher Scientific, Hampton, NH) suspended within a 50 mL centrifuge tube to remove any remaining particulates. The flow-through from this filter was then passed through a 35 µm cell strainer into a flow tube (Corning, Corning, NY). Flow tubes were centrifuged at 250 rcf for 5 min, the supernatant was removed, and the cell pellets were resuspended in 50 µL Flow Cytometry Staining Buffer (Invitrogen). Cell concentration was quantified using a Countess II FL (Invitrogen) and each sample was diluted to 2*10^7^ cells per mL. Finally, 50 µL of each diluted sample added to 5 mL tubes (Eppendorf, Hamburg, DE), resulting in 1 million cells per flow sample.

Harvested spleens were placed into a prewetted 100 µm cell strainer (Fisher Scientific, Hampton, NH) in a 50 mL centrifuge tube and mashed aggressively with the back of a plunger from a sterile 10 mL syringe (Becton Dickinson) with repeated rinsing using sterile Hanks’ Balanced Salt Solution (HBSS). Flow-through from this filter was then pipetted into a 70 µm cell strainer into a new 50 mL centrifuge tube; this flow-through was then pipetted through a 35 µm cell strainer cap into a flow tube. The samples were centrifuged at 250 rcf for 5 min at 4 °C, the supernatant was removed, and cell pellets were resuspended in 1 mL ice-cold ACK lysis buffer (Quality Biological, Gaithersburg, MD). Red blood cells were lysed for 30 sec before the solution was pipetted into 10 mL cold PBS. Tubes were centrifuged at 250 rcf for 5 min at 4 °C, the supernatant was removed, and pellets were resuspended in 500 µL flow buffer. Cell densities were calculated using the Countess II FL, and samples were diluted to 2*10^7^ cells per mL; 50 µL of each diluted solution was then added to 5 mL tubes to constitute 1 million cells per flow sample.

Once normalized by cell count, both spleen and LN flow samples were processed identically, with complete flow panels shown in Table S2. Each tube of cells then received 2 µL of Fc block (BioLegend, San Diego, CA), 10 µL of Brilliant Stain Buffer (Becton Dickinson), and 28 µL of flow buffer. Additionally, each tube in both panels received 2.5 µL of anti-mouse H-2K^b^-SIINFEKL antibody conjugated to allophycocyanin; we previously confirmed that SKH1-Elite mice express the H-2K^b^ haplotype of MHC I.^28^ All tubes were incubated on ice in the dark for 45 min. The remaining antibodies were added and the samples were incubated on ice in the dark for an additional 30 min. Next, 1 mL cold PBS was added to each tube and samples were centrifuged at 200 rcf at 4 °C for 7 min. Supernatant was removed and cell pellets were resuspended in 1 mL PBS containing a LIVE/DEAD stain (Table S2); samples were incubated on ice in the dark for 30 min. Finally, cells were centrifuged at 200 rcf at 4 °C for 7 min, the supernatant was removed, and cell pellets were resuspended in 1 mL flow buffer. The solutions were passed through 35 µm cell strainer caps into flow tubes and analyzed by flow cytometry; spleen samples were measured using a Sony SA3800 Cytometer and LN samples using a Sony MA900 Cell Sorter (Sony Biotechnology, San Jose, CA).

### Statistics

All group comparisons were calculated by ordinary one-way ANOVA with Tukey’s multiple comparison test to compare the performance of the PEG-OVA + “Click” group to all other groups. All statistical analyses were performed in GraphPad Prism 10. Statistical significance in all figures is denoted with asterisks as follows: * = p < 0.05; ** = p < 0.01; *** = p < 0.001; **** = p < 0.0001. Error bars indicate standard deviation in bar graphs and standard error of the mean in line graphs.

## Results and Discussion

### Click-functionalized PEG crosslinks rapidly and irreversibly, entrapping bound OVA

To prepare materials capable of bioorthogonal crosslinking, we first synthesized Tz- and TCO-terminated 8-arm PEG in house; TCO-PEG was additionally functionalized with Cy5, and Tz-PEG with DyLight 755-labeled OVA. We calculated a total of 7.0 Tz per PEG molecule in the Tz-PEG material via UV-VIS spectroscopy. After conjugation with OVA-DyLight 755 and purification via SEC, the final material contained 1:4.04 OVA:PEG as determined via fluorescence measurements. Similarly, TCO-PEG was determined to have 6.8 TCO per PEG molecule by UV-VIS and 0.10:1 Cy5:PEG by fluorescence measurements. Endotoxin levels were at or below the limit of detection of 0.15 EU/mg for all materials, below advised limits for preclinical research.^29^ Both PEG materials had a hydrodynamic diameter of ∼10 nm as determined via DLS, which is within the size range that preferentially passes through the lymphatic vessels instead of intravasating into blood capillaries from the subcutaneous space (Figure S2).

We then measured the degradation rate of click hydrogels by performing an in vitro release assay using blind well chambers. The crosslinking group contained equal parts Tz-PEG-OVA and TCO-PEG, while the non-crosslinking control group contained equal parts Tz-PEG-OVA and Tz-PEG. The high rate at which TCO and Tz react allowed for visible gelation at high concentrations (7.5% w/v) immediately after pipetting together. The quantity of OVA released was assessed by measuring the fluorescence intensity of DyLight 755 removed from the top chamber of a blind well (Figure S3). We found that OVA release was completed within 2 days in the non-crosslinking materials, whereas only 21 ± 1% of OVA had been released from the crosslinked hydrogels. Over the next 11 days, minimal amounts of OVA were released from the crosslinking group, demonstrating that these materials crosslink and do not meaningfully degrade via hydrolysis on this timescale *in vitro*.

### Vaccination with crosslinkable components increases antigen presence and retention in the draining LN

To determine if the *in situ* crosslinking of vaccine-loaded hydrogels within LNs improves antigen trafficking and/or retention of antigen in the LN, SKH1-Elite mice were vaccinated using the complete LN-targeting system, henceforth called PEG-OVA + “Click,” control groups that progressively remove aspects of the system, and the traditional alum adjuvant as a positive control (Figure 1A). At 6 h post-vaccination, all injected components were visible within the inguinal LN (InLN) by IVIS imaging (Figure 1B). OVA signal in the InLN was quantified over one week by manual positioning of an ROI over the visible LN spot (Figure 2A). The peak intensity for all groups occurred at 6 h, and the OVA-DyLight 755 signal was highest in the PEG-OVA + “Click” group, which was statistically significantly greater than all but the PEG + Free OVA + “Click” group (Figure 2B,C). This rapid movement to the LN suggests that OVA traveled through passive lymphatic flow rather than by cell-mediated uptake, which normally requires 18-24 h.^30^ The OVA signal in all groups exponentially decayed beginning after 6 h. The area under the curve calculated in radiant efficiency over time showed a significant increase in the PEG-OVA + “Click” group over all other groups (Figure 2C).

**Figure 1:**
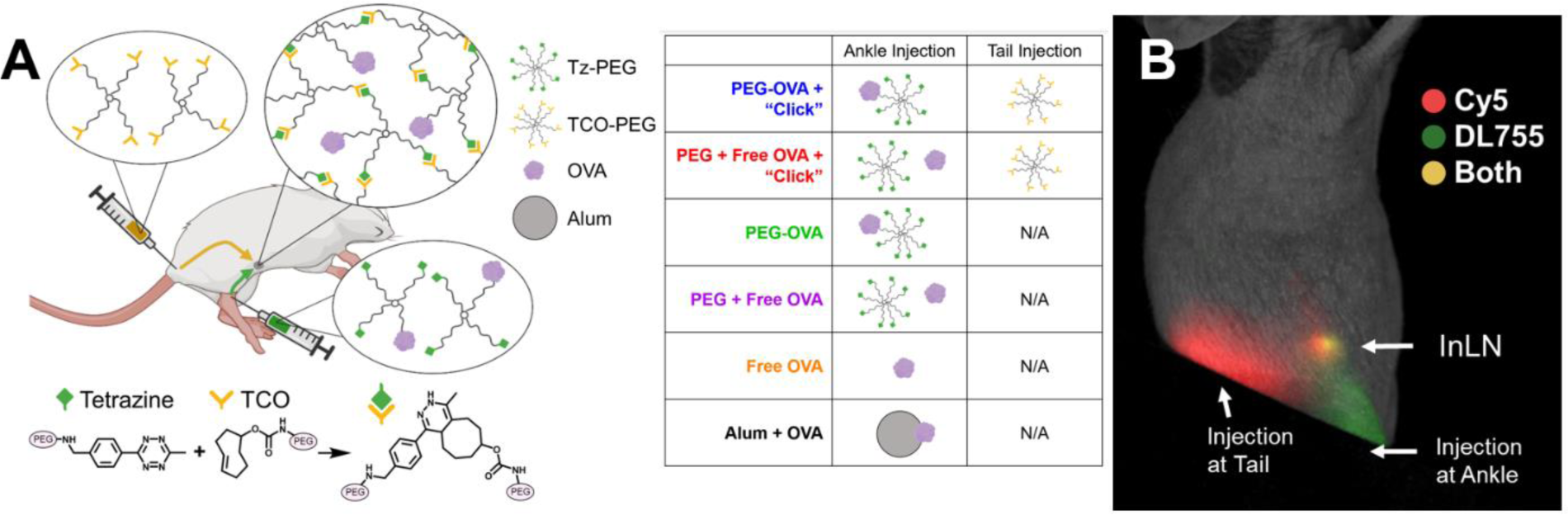
The lymph node crosslinking platform. **(A)** Illustration of the platform mechanism. The two components are injected into either the tail base or ankle, which both preferentially flow to the inguinal lymph node (InLN) where they rapidly crosslink. The table provides a key of the components administered in each of the groups used throughout subsequent experiments; group names are color-coded for consistency across all further figures. **(B)** Spectrally unmixed IVIS image of a mouse in the PEG-OVA + “Click” group 6 h after vaccination. Note: A black index card was placed over the injection sites to allow for easier visualization of the InLN. Cy5 was conjugated onto PEG-TCO and DyLight 755 was conjugated to OVA.

**Figure 2:**
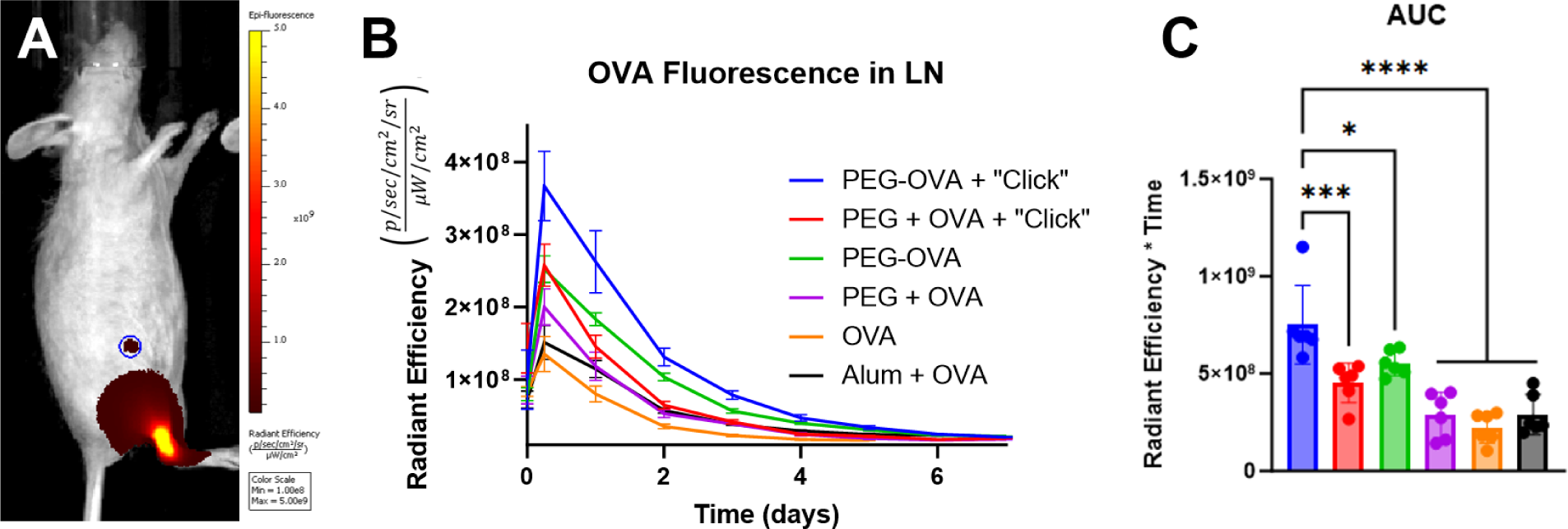
Ovalbumin collection and retention in the LN. **(A)** Representative IVIS image showing DyLight 755-OVA signal from the PEG-OVA + “Click” group in the InLN 24 hours after vaccination. The blue circle represents the region of interest used to calculate **(B)**, OVA presence and retention in the LN over time. **(C)** Area-under-the-curve calculations of data in (B). Note: The legend in (B) also applies to (C).

The higher intensity and prolonged duration of fluorescence in the LN in the PEG-OVA + “Click” group compared to their non-crosslinking controls implies that the ability to crosslink within the LN extends OVA residence time, thereby leading to a greater accumulation of OVA. No notable differences in the drainage of OVA from the injection site were noted in any groups other than Alum + OVA (Figure S4), which is expected since alum-adsorbed antigens are known to slowly desorb and thereby prolong release. The elevation of signal in the PEG + Free OVA + “Click” group at early timepoints suggests that OVA may briefly become entrapped when co-delivered with unconjugated PEG as the PEG crosslinks with the complementary click component. Similarly, the elevation of signal in the PEG-OVA group suggests that conjugation of PEG to OVA may improve trafficking of the antigen into the lymphatics. The synergism of improved trafficking and *in situ* crosslinking then produces the highest overall signal in the PEG-OVA + “Click” group.

### Injected components are visible within the LN after 24 hours by immunofluorescence

To better understand how the system functions at the cellular level, draining InLNs were harvested for immunofluorescence analysis. While OVA signal was visible in cryosections from all groups in the medulla, just upstream of the efferent lymphatics, or the “exit” of the LN (Figure 3A), sections from the crosslinking groups (PEG-OVA + “Click” and PEG + Free OVA + “Click”) were more likely to show OVA near the “entrance,” on the afferent end of the LN (Figure 3B). In these sections, both the OVA and the complementary TCO-PEG component were colocalized with CD169^+^ macrophages, with some of the PEG material appearing to additionally be present within macrophages (Figure 3B, arrowheads). Representative sections from the other treatment groups can be seen in Figure S5; all non-crosslinking groups (PEG-OVA, PEG + Free OVA, Free OVA, and Alum) appeared similar.

**Figure 3:**
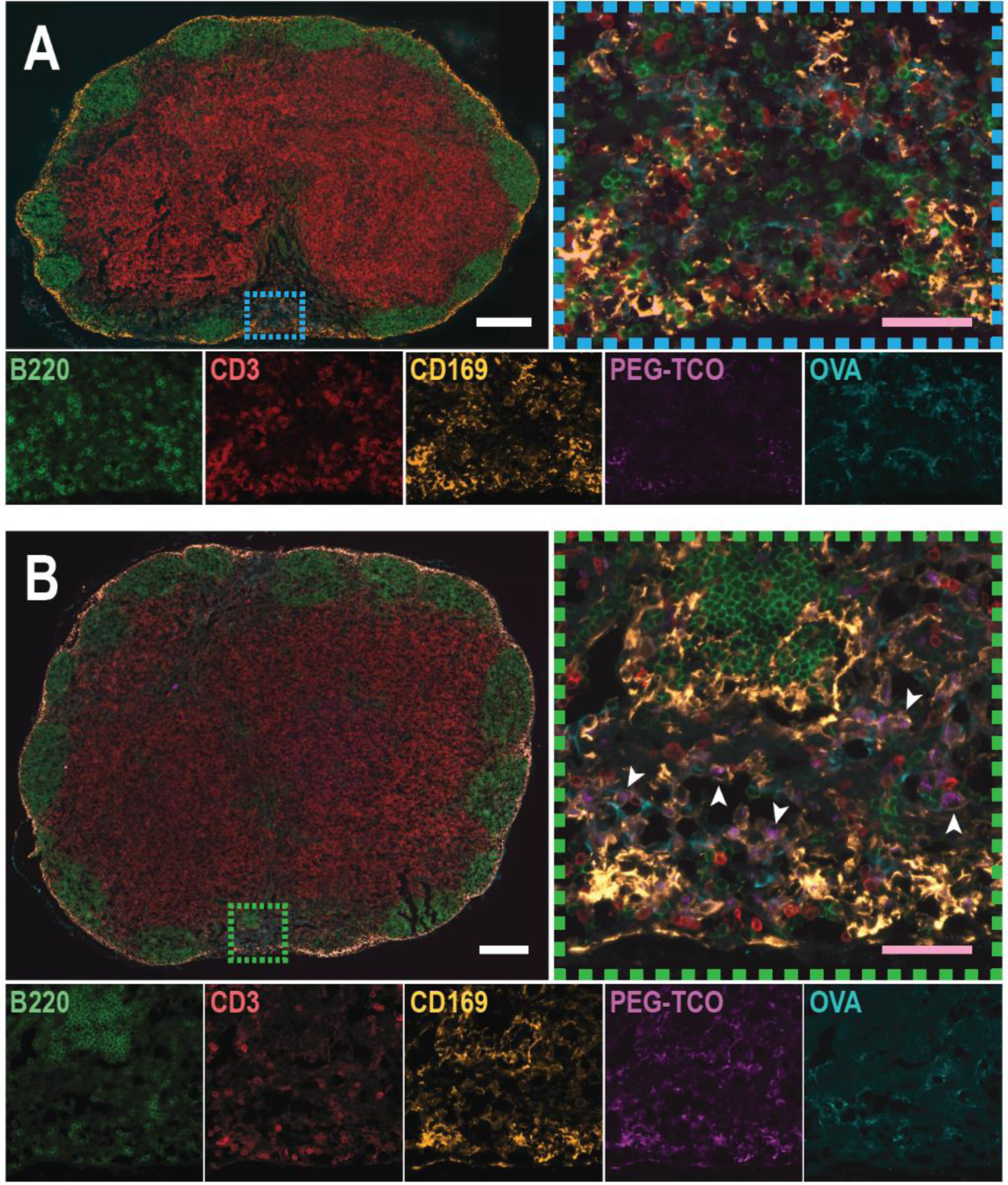
Immunofluorescence staining of targeted inguinal LNs 24 h after material administration. **(A)** Lymph node section from the Free OVA group, also representative of sections from the PEG-OVA, PEG + Free OVA, and Alum + OVA groups. The blue dashed box shows the area zoomed in on the right, showing the presence of OVA in the medulla. Individual fluorescence channels for the zoomed image are shown below: Alexa Fluor 488 (B220, B cells), Spark YG 570 (CD3, T cells), Alexa Fluor 594 (CD169, macrophages), Cy5 (PEG-TCO), and DyLight 755 (OVA, amplified with anti-OVA antibody). Note: some bleedover was seen in the Cy5 channel of Alexa Fluor 594; when merged, the dimmer bleedthrough into Cy5 is suppressed under Alexa Fluor 594, allowing true Cy5 signal to be visible. **(B)** LN section from the PEG-OVA + “Click” group, also representative of sections from the PEG + Free OVA + “Click” group. The green dashed box shows the area zoomed in on the right, demonstrating the colocalization of PEG-TCO and OVA in the medulla and the apparent encapsulation of PEG-TCO by CD169^+^ macrophages (white arrowheads). White scale bars denote 250 µm, pink scale bars denote 50 µm.

CD169^+^ macrophages, sometimes called “gatekeepers,” are the first cells to capture antigens in the LN and are understood to interact with resident DCs, which in turn activate T cells.^31^ CD169 is expressed on both subcapsular sinus macrophages (SSMs) and medullary sinus macrophages (MSMs); we presume that the CD169^+^ cells encapsulating our PEG materials are MSMs due to both their location outside of the subcapsular sinus and previous studies showing that SSMs are poorly endocytic relative to MSMs.^32,33^ In addition, antigens delivered by passive lymphatic flow are thought to be primarily captured by MSMs.^33^ Although it appears that the Cy5-conjugated TCO-PEG is the only material within the MSMs, the physical size of the Cy5 particulates (2-8 µm in diameter) suggests that they are assemblies of hundreds to thousands of 10 nm PEG molecules. We hypothesized that this was a result of crosslinking with Tz-PEG, which was not fluorescently labeled. To test this hypothesis, we removed Tz-PEG from the materials administered in order to prevent click chemistry-mediated assembly and injected mice with TCO-PEG component in the tail base and free OVA in the ankle. We found limited Cy5 signal, indicative of TCO-PEG, visible in the lymph nodes and no visible pockets of TCO-PEG-Cy5 in MSMs, suggesting that these microscale particulates are a result of TCO-PEG crosslinking with Tz-PEG (Figure S6).

### The ability to crosslink in the LN improves humoral immunity

To determine how increased persistence, retention, and cell uptake affected the humoral immune response, we evaluated antigen-specific antibody titers over 14 weeks following vaccination (Figure 4A,B). At the week 14 endpoint, the PEG-OVA + “Click” group had significantly higher IgG titers than the three non-adjuvanted non-crosslinking groups, exhibiting a 256-fold increase in geometric mean over the Free OVA control. The PEG + Free OVA + “Click” group had average titers 12-fold lower than the PEG-OVA + “Click” group, though this difference was not statistically significant. Interestingly, despite producing the second-highest OVA fluorescence profile in the LN, PEG-OVA produced antibody titers equivalent to the OVA control and on average 80-fold lower than PEG-OVA + “Click.” These findings suggest that, while conjugation to multi-arm PEG enhanced OVA trafficking to the lymphatics, this improvement had no discernable effect on its humoral immunity. Meanwhile, the elevation of titers in the PEG + Free OVA + “Click” group relative to the material-matched PEG + Free OVA group implies that there is an immunological benefit to crosslinking within the draining LN. Further, crosslinking, rather than general material adjuvancy, appears to be responsible for a majority of the immunological benefit since the material-matched PEG + Free OVA group produced titers that, like the unadjuvanted OVA control treatment, were below the limit of detection. The clinically common adjuvant, alum, which is known to primarily promote humoral immunity, produced very high OVA-specific IgG titers that were significantly greater than all non-adjuvanted groups. Together, these data show that this platform produces an improvement in humoral immune response over dose-matched non-crosslinking controls without employing a pathogen-mimicking adjuvant.

**Figure 4:**
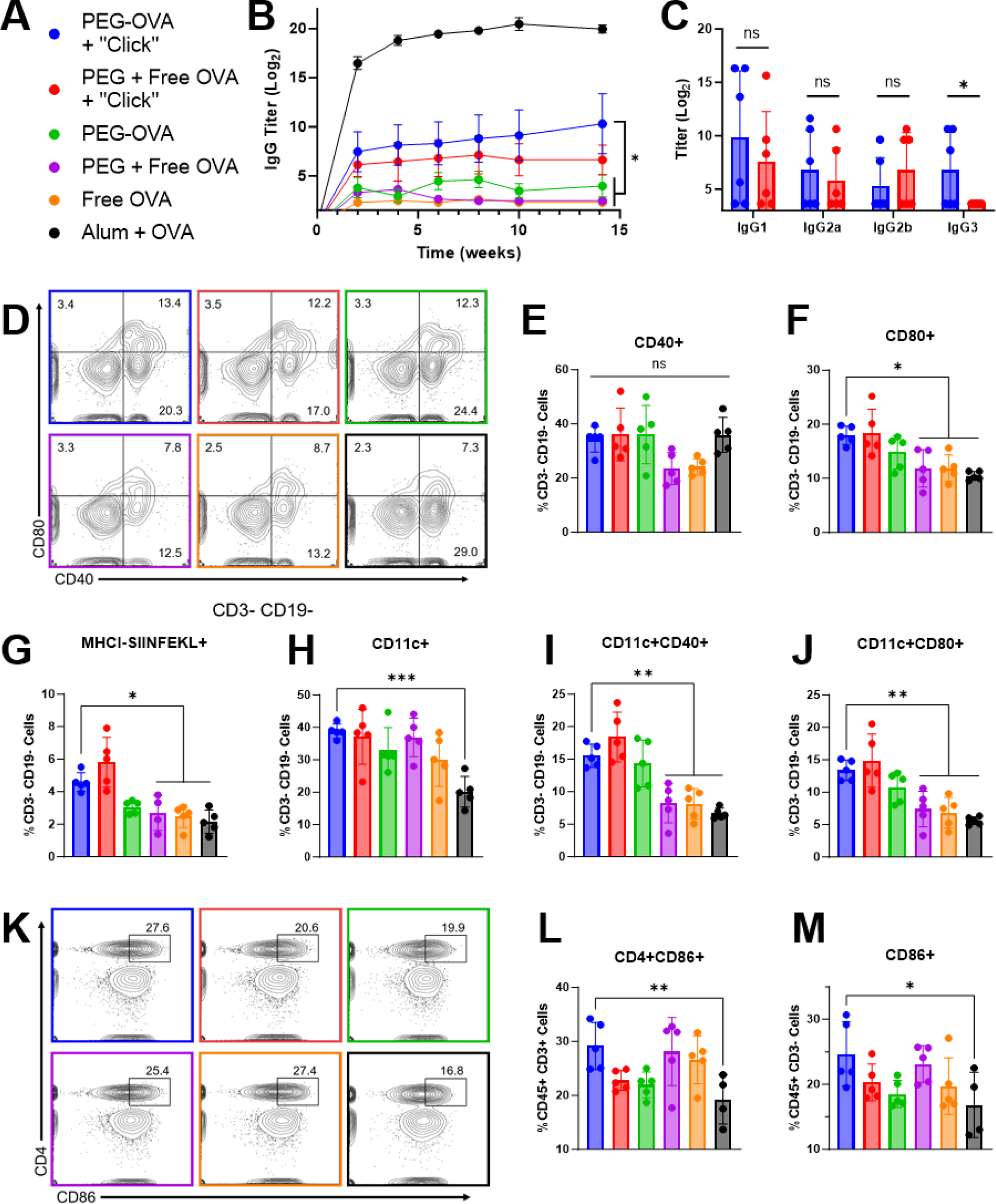
Immunological characterization of the intra-LN crosslinking platform. **(A)** Legend applies to all panels in this figure. **(B)** Anti-OVA IgG titers measured over 14 weeks for all groups, n=6. PEG-OVA + “Click” is statistically greater than PEG-OVA, PEG+ Free OVA, and Free OVA at week 14. Alum is statistically greater than all other groups at week 14. Error bars represent SEM. **(C)** Anti-OVA IgG subclasses measured for the two crosslinking groups, n=6. PEG-OVA, PEG + Free OVA, and Free OVA are not presented as all were below the limit of detection. **(D-J)** Flow cytometry of cells from targeted lymph nodes, n=5. **(D)** Representative plots of non-B/T lymphocytes expressing the activation markers CD80 and CD40. A graphical representation of each of these markers is shown in **(E)** and **(F). (G)** Expression of MHC I presenting the OVA peptide SIINFEKL in non-B/T lymphocytes. **(H)** Percent of non-B/T cells expressing the dendritic cell marker CD11c. **(I)** Percent of non-B/T cells co-expressing CD11c and CD40. **(J)** Percent of non-B/T cells co-expressing CD11c and CD80. **(K-M)** Flow cytometry of spleens, n=4-5. **(K)** Representative plots of T cells expressing CD4 and the activation marker CD86, with the graphical representation shown in **(L)**. **(M)** Percent of non-T lymphocytes expressing CD86.

The concentration of IgG subclasses in blood was measured 10 weeks after vaccination for both crosslinking groups, which were the only adjuvant-free formulations that elicited total anti-OVA IgG titers above the limit of detection (Figure 4C). Both groups had statistically similar titers for all subclasses except IgG3, for which the PEG-OVA + “Click” group was significantly greater. Mouse IgG3, which is not homologous to human IgG3, is T cell-independent and, when bound to a protein antigen, has been found to colocalize with follicular dendritic cells and further enhance the generation of antigen-specific antibodies.^34^ While its precise receptor on macrophages is still being investigated,^35^ murine IgG3 has also been hypothesized to multimerize to effectively increase its affinity and avidity.^36^ The alum-adjuvanted group produced significantly greater titers for all subclasses (Figure S7), which was expected based on the high total IgG titers obtained in that group.

### Improved activation of antigen-presenting in lymph nodes and spleens compared to alum

The cellular immune response to vaccination with *in situ* clicking hydrogels was assessed by performing flow cytometry on cells isolated from InLNs and spleens one week after vaccination. Flow cytometry of cells from draining LNs showed that vaccination with PEG-OVA + “Click” significantly increased CD80-activated non-T and non-B cells, identified as CD3^-^CD19^-^, compared to PEG + Free OVA, Free OVA, and Alum + OVA, with many of those cells also expressing CD40 (Figure 4D-F). LN-resident cells that are not T or B cells are likely to be APCs—most commonly macrophages and DCs. CD80 expression on the surface of APCs provides co-stimulation for T cells by binding to CD28, inducing proliferation and differentiation; without CD28 ligation, T cell receptor binding induces anergy or apoptosis.^37^ Expression of CD80 is also commonly used to identify M1-polarized, “classically-activated” macrophages prevalent in infections. We observed a significant increase in cells in this non-B/T subset expressing MHC I presenting SIINFEKL, a peptide portion of OVA, in the PEG-OVA + “Click” group relative to PEG + Free OVA, Free OVA, and Alum + OVA and nearly significant over PEG-OVA (p=0.054) (Figure 4G). This signifies cross-presentation of antigen by MHC I, which is specifically involved in the activation of CD8^+^ T cells.^38^

Probing this non-B/T cell group further, we found a significant increase in the percent of DCs, using CD11c as a marker, in the PEG-OVA + “Click” group compared to Alum + OVA group (Figure 4H). There was also a significant increase in activation of DCs, as evidenced by a concurrent increase in CD40 and CD80 expression in the PEG-OVA + “Click” group relative to the PEG + Free OVA, Free OVA, and Alum + OVA groups (Figure 3I,J). The expression and upregulation of CD40 on DCs has been shown to improve their antigen presentation and help trigger other co-stimulatory molecules to activate T cells.^39,40^ In all aforementioned assessments of antigen-presenting cells, the PEG + Free OVA + “Click” group and the PEG-OVA group performed comparably to the PEG-OVA + “Click” group (Figure 4D-J). We hypothesize that this is due to the increased LN residence time of OVA in these groups over the PEG + Free OVA, Free OVA, and Alum + OVA groups (Figure 2B,C).

Analysis of CD3^+^ T cells and CD19^+^ B cells in the LN did not show any significant differences within the groups at this timepoint. We also did not find any significant differences between draining and contralateral InLNs within each mouse in any measured property. The non-targeted LNs showed similar levels of all markers, including DC activation, as the draining LN from the same mouse (data not shown). These similarities suggest a more systemic response by the immune system at 1 week after vaccination by this platform rather than evoking a response that is isolated to only the draining LN, which is useful for vaccines against infectious diseases.^41^

Flow cytometry of splenocytes isolated from the same mice showed a significant increase in non-T lymphocytes, identified as CD45^+^CD3^-^, expressing the activation marker CD86 in the PEG-OVA + “Click” group over the Alum + OVA group (Figure 4K,L). CD86, like CD80, is a costimulatory molecule expressed on APCs that binds to CD28 on T cells. As this group contains all professional APCs, this increase may be correlated to the enhanced activation of APCs seen in the LNs, although no significant difference is seen from PEG + Free OVA or Free OVA as was detected in the LNs (Figure 4F). The PEG-OVA + “Click” group also had elevated levels of activated helper CD4^+^ T cells expressing CD86 compared to the Alum + OVA group (Figure 3M); although CD86 is predominantly expressed on APCs, its expression on T cells has been correlated to an activated effector function and naïve T cell priming.^42,43^

## Conclusions

Together, these data show that the crosslinking of PEG molecules functionalized with complementary click groups in the draining LN enhances the immune response to antigens compared to material-matched controls. This platform exerts this effect despite the fact that PEG resists recognition by the immune system and has been shown to have no adjuvant effect in mice.^44^ This work shows that the covalent attachment of OVA onto multi-arm PEG (PEG-OVA) improves trafficking to and retention within the InLN from a subcutaneous ankle injection. This formulation increases APC activation within the LN but has a minimal effect on the antibody response. When soluble OVA is co-delivered with PEG that is capable of forming crosslinked microgels within the LN via a click reaction (PEG + Free OVA + “Click”), it similarly increases LN APC activation but, unlike the uncrosslinkable materials, also increases OVA-specific IgG titers approximately 20-fold compared to the Free OVA control group. Covalently attaching OVA onto PEG in a click-reaction capable system (PEG-OVA + “Click”) further improved antibody titers, increasing them 80-fold compared to the PEG-OVA control and 256-fold compared to the Free OVA control while increasing LN APC activation.

Alum is known to be an excellent adjuvant for conferring humoral immunity but is far from optimal at enhancing cellular immunity. A vaccination platform that uses passive lymphatic drainage to access the lymph node presents several advantages over peripheral uptake and trafficking by tissue-resident DCs—the mechanism favored in response to alum—especially in the context of cellular immunity. Systems that employ passive lymphatic drainage produce a higher amount of antigen-presenting DCs,^2^ display antigen to T cells in the LN more quickly,^45^ and increases the likelihood that the antigen remains in its native conformation.^46^ By increasing antigen accumulation and retention in the LN, our platform improves DC activation and MHC I presentation of antigen compared to alum without help from a traditional adjuvant. This platform also can also be readily customized through reactions with its TCO and Tz end groups, allowing for the addition or substitution of other antigens or adjuvants, which makes it an attractive system for a wide variety of vaccines requiring different types of immune response.

## Acknowledgements

The authors thank Lynna Baryakova and Mei-Li Laracuente for their advice and aid. This work was supported by National Institute of Health grants K22AI146215 and R35GM143101, the Cancer Prevention and Research Institute of Texas grant RR190056, and start-up funds from Rice University. The graphic in Figure 1A was made using biorender.com.

