## Supplemental Information for "Intra-Lymph Node Crosslinking of Antigen-Bearing Polymers Enhances Humoral Immunity and Dendritic Cell Activation"

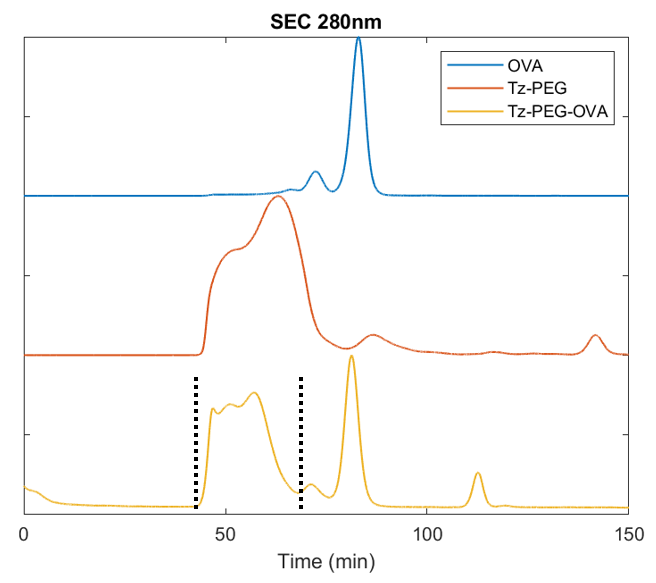

**Figure S1**: Size Exclusion Chromatography (SEC) chromatograms at 280 nm of OVA, PEG, and PEG-OVA to illustrate the purification method. The duration of material collection of PEG in the Tz-PEG-OVA group is shown by the dotted lines. All chromatograms were scaled to the same maximum intensity.

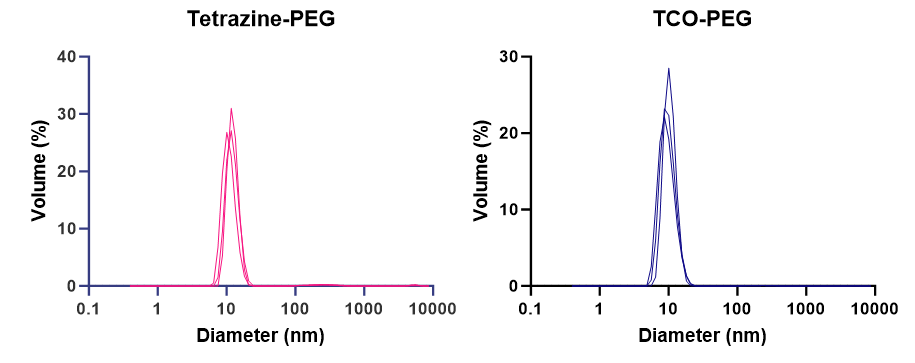

**Figure S2**: Dynamic Light Scattering (DLS) volume-weighted measurements of Tz-PEG and TCO-PEG, showing maximum diameters around 10 nm.

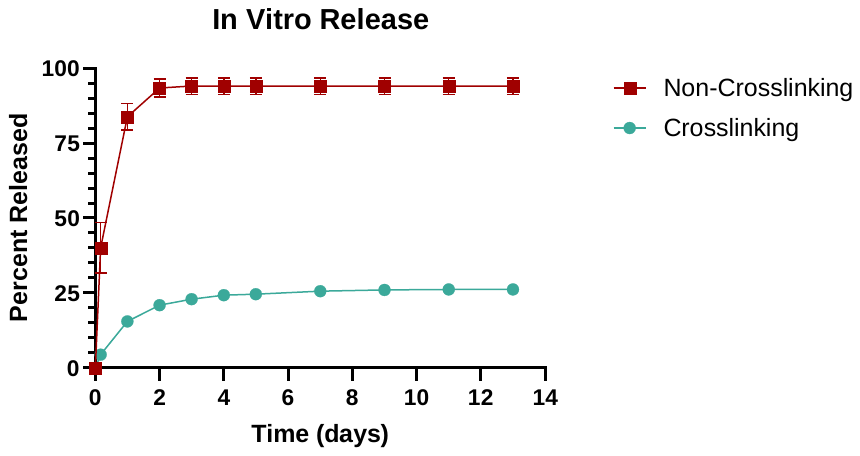

**Figure S3**: *In vitro* release of OVA using blind well chambers (n=3). The crosslinking group used an equal mixture of TCO-PEG and Tz-PEG-OVA, while the non-crosslinking group used an equal mixture of Tz-PEG and Tz-PEG-OVA.

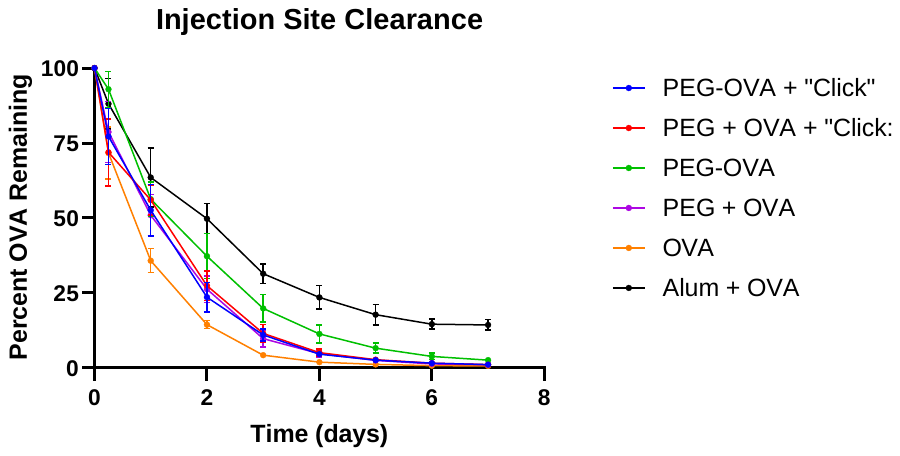

**Figure S4**: Clearance of OVA from the injection site in the ankle, normalized to post-injection values (n=6).

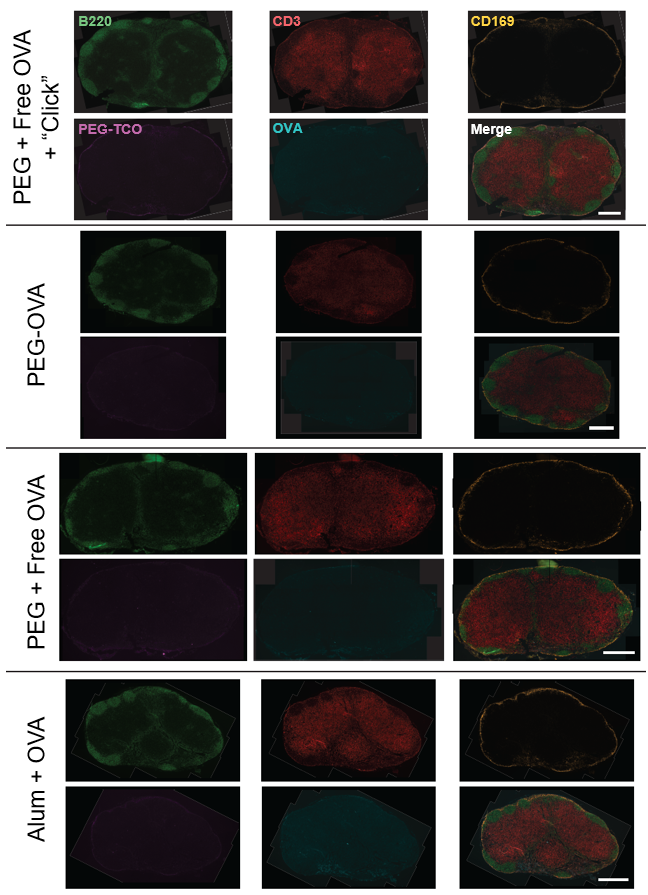

**Figure S5**: Immunofluorescence staining of lymph node sections. Scale bars are 500 µm.

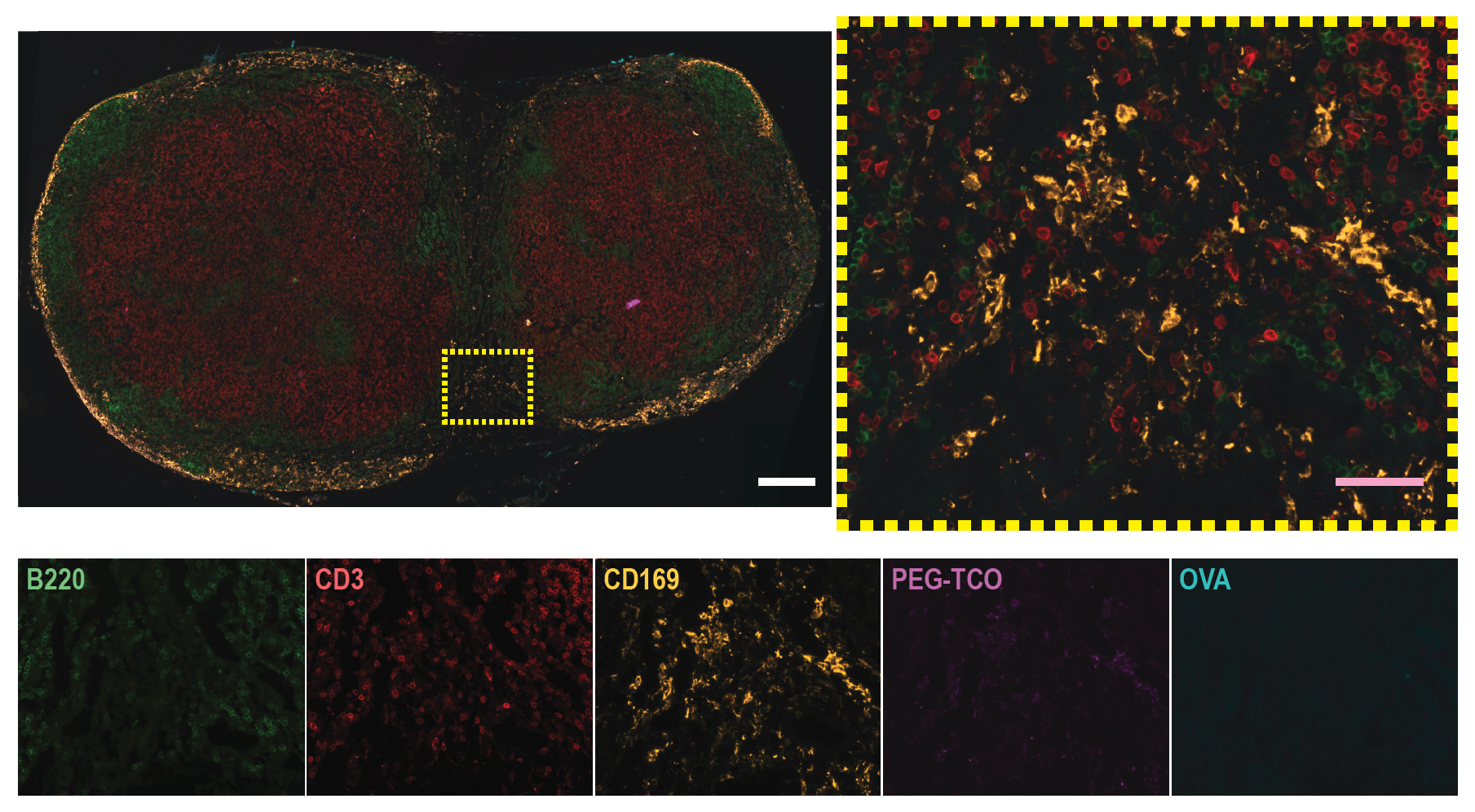

**Figure S6**: Immunofluorescence staining of lymph node sections from a control group receiving a tail base injection of TCO-PEG and an ankle injection of free OVA. No microscale particulates of TCO-PEG are visible in CD169^+^ macrophages. Individual fluorescence channels for the zoomed image are shown below: Alexa Fluor 488 (B220, B cells), Spark YG 570 (CD3, T cells), Alexa Fluor 594 (CD169, macrophages), Cy5 (PEG-TCO), and DyLight 755 (OVA, amplified with anti-OVA antibody). Note: some bleedover was seen in the Cy5 channel of Alexa Fluor 594; when merged, the dimmer bleedthrough into Cy5 is suppressed under Alexa Fluor 594, allowing true Cy5 signal to be visible. White scale bars denote 250 µm, pink scale bars denote 50 µm.

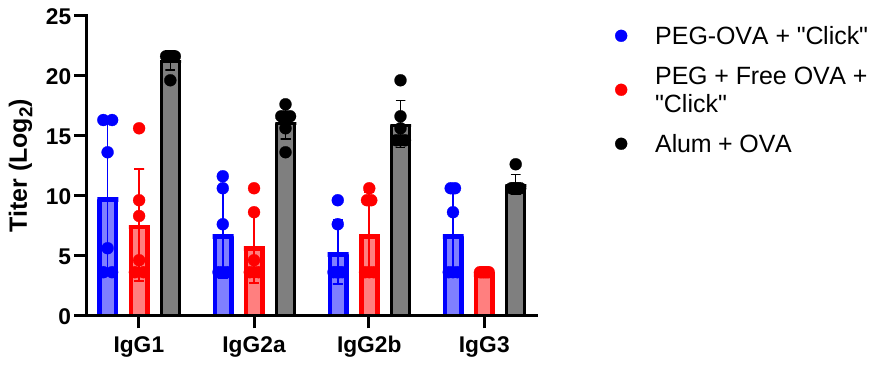

**Figure S7**: Antibody subclass analysis (n=6). Alum + OVA is significantly greater than the PEG-OVA + “Click” and PEG + Free OVA + “Click” groups in all subclasses.

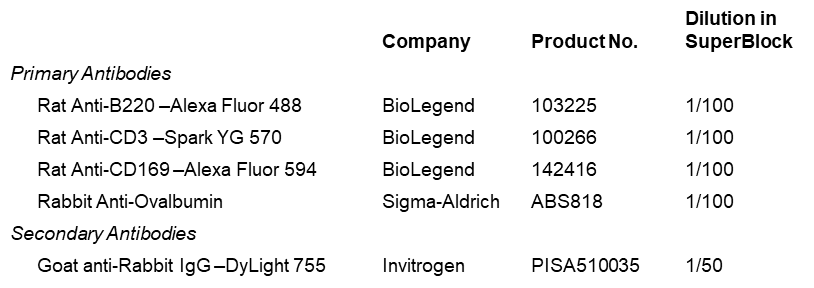

**Table S1**: Antibodies used for immunofluorescence staining of lymph node sections.

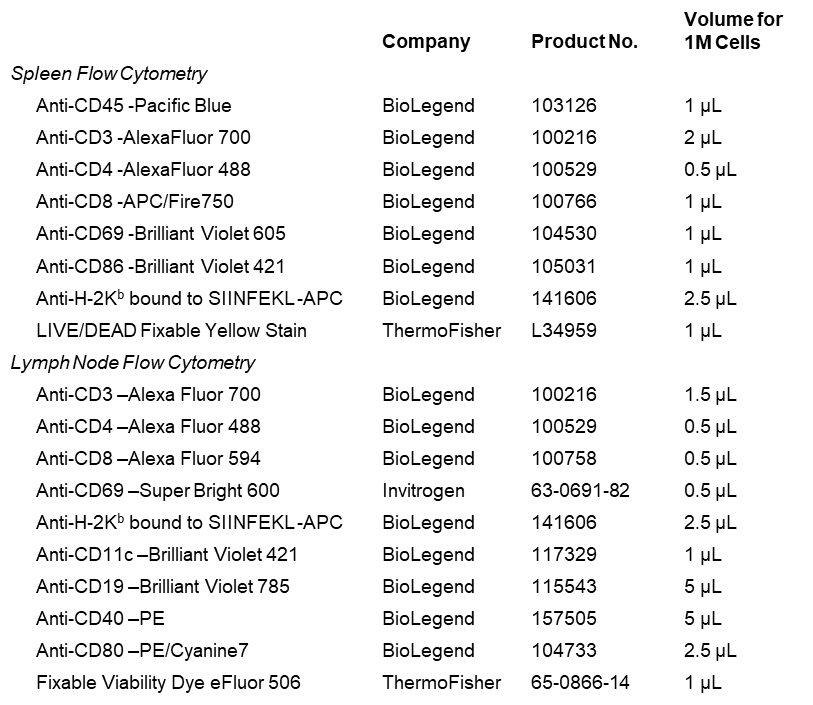

**Table S2**: Antibodies and stains used for flow cytometry.
